# Cross-reactivity of a rice NLR immune receptor to distinct effectors from the blast pathogen leads to partial disease resistance

**DOI:** 10.1101/530675

**Authors:** Freya A. Varden, Hiromasa Saitoh, Kae Yoshino, Marina Franceschetti, Sophien Kamoun, Ryohei Terauchi, Mark J. Banfield

## Abstract

Unconventional integrated domains in plant intracellular immune receptors (NLRs) can directly bind translocated pathogen effector proteins to initiate an immune response. The rice immune receptor pairs Pik-1/Pik-2 and RGA5/RGA4 both use integrated heavy metal-associated (HMA) domains to bind the *Magnaporthe oryzae* effectors AVR-Pik and AVR-Pia, respectively. These effectors both belong to the MAX effector family and share a core structural fold, despite being divergent in sequence. How integrated domains maintain specificity of recognition, even for structurally similar effectors, has implications for understanding plant immune receptor evolution and function. Here we show that the rice NLR pair Pikp-1/Pikp-2 triggers an immune response leading to partial disease resistance towards the “mismatched” effector AVR-Pia in planta, and that the Pikp-HMA domain binds AVR-Pia in vitro. The HMA domain from another Pik-1 allele, Pikm, is unable to bind AVR-Pia, and does not trigger a response in plants. The crystal structure of Pikp-HMA bound to AVR-Pia reveals a different binding interface compared to AVR-Pik effectors, suggesting plasticity in integrated domain/effector interactions. This work shows how a single NLR can bait multiple pathogen effectors via an integrated domain, and may enable engineering immune receptors with extended disease resistance profiles.

## INTRODUCTION

When plants encounter biotic stresses, they respond rapidly to defend themselves against attack. Microbial pathogens translocate effector proteins inside host cells to undermine plant immunity and promote pathogen growth and proliferation (1). To detect these effectors, plants have developed intracellular immune receptors, many of which are of the NLR (nucleotide-binding leucine-rich repeat) class (2). The hallmark feature of NLR-mediated immunity is the hypersensitive response (HR), a programmed cell death around the site of infection that helps to isolate and halt the spread of the pathogen (3).

NLRs recognise effector proteins via different mechanisms, including by direct or indirect binding (4,5). Some NLRs function in pairs, with one receptor responsible for recognising the effector (referred to as the sensor), and one responsible for translating the recognition into a signalling response (the helper) (6). One mechanism to evolve direct binding has been for NLRs to integrate an unconventional domain into the protein architecture (7,8), with this domain thought to be derived from the virulence-associated host target of the effector. Once integrated, these domains may adapt to recognise effectors (and different effector alleles). Their widespread distribution in NLRs from diverse plant species suggests this is an ancient mechanism for evolving effector recognition (9,10).

Two paired rice NLR immune receptors are known that contain an integrated heavy metal-associated (HMA) domain, Pik-1/Pik-2 and RGA5/RGA4. In Pik, this domain is integrated between the coiled-coil (CC) and nucleotide-binding (NB-ARC) domains of Pik-1 (11,12), whereas in the RGA pair the HMA domain is found at the C-terminus of RGA5 (13). Both these pairs of immune receptors recognise effectors from the blast fungus *Magnaporthe oryzae*, a global threat to rice production causing loss of up to a third of the total annual harvest of this crop (14-16).

*M. oryzae* secretes a large repertoire of effector proteins and many of these, including the structurally characterised AVR-Pizt, AVR-Pia, AVR-Pik, AVR1-CO39 and AVR-Pib (11,17-19), share a conserved structure comprising a six stranded β-sandwich known as the MAX (*Magnaporthe* Avrs and ToxB-like) fold (18,20). Therefore, despite being sequence-unrelated, these effectors are all similar in overall shape.

The Pik-1/Pik-2 NLR pair recognise the *M. oryzae* effector AVR-Pik (21), and both the NLRs and effectors are found as allelic series in natural populations (22). Direct interaction between the Pik-HMA domain and AVR-Pik is required for triggering an immune response to the effector (11). At the sequence level, the allelic Pikp (23) and Pikm (24) pair differ mainly in their polymorphic HMA domains (12) and this underpins different recognition specificities for different AVR-Pik alleles; Pikp is only able to recognise the effector variant AVR-PikD, whereas Pikm can recognise AVR-PikD and other additional AVR-Pik variants. The AVR-PikC effector variant is currently unrecognised by any Pik NLR.

The RGA5/RGA4 NLR pair responds to the *M. oryzae* effectors AVR-Pia (25) and AVR1-CO39 (13). Both AVR-Pia and AVR1-CO39 physically interact with RGA5-HMA and this interaction is required for triggering resistance (13,26).

Despite similarities in the Pik-1/Pik-2 and RGA5/RGA4 systems, their mechanisms of activation are different. The Pik-1/Pik-2 pair appear to use a cooperative mechanism, where effector recognition by the HMA in the sensor NLR Pik-1 requires the helper NLR Pik-2 to initiate signalling, but Pik-2 cannot signal on its own. Contrastingly, the RGA5/RGA4 pair functions via negative regulation, where recognition of the effector through RGA5-HMA derepresses signalling by RGA4 (27,28). However, details of the NLR interactions and the resultant downstream signalling remain to be understood.

The interface between AVR-Pik effectors and the HMA domain of both Pikp and Pikm has been extensively studied and structurally characterised (11,12). Recently, the structure of AVR1-CO39 in complex with the HMA domain of RGA5 was also elucidated (29), and revealed that the HMA/effector interface was substantially different compared to the Pik NLR pairs. This has raised intriguing questions concerning how structurally similar but sequence divergent HMA domains distinguish between structurally similar but sequence divergent pathogen effectors.

Here we reveal that Pikp is able to trigger partial disease resistance to the “mis-matched” effector AVR-Pia in rice, and elicits a weak cell death response in *N. benthamiana*. Pikp-HMA binds AVR-Pia in vitro, at the RGA5/AVR1-CO39-like interface, rather than the Pik/AVR-Pik-like interface. This structural understanding of effector cross-reactivity in the Pik/RGA systems provides insights into the evolution and function of integrated HMA domains in NLRs. It also hints at the potential to engineer the HMA of Pikp to respond robustly to both AVR-PikD and AVR-Pia at the different interfaces.

## RESULTS

### Co-expression of Pikp/AVR-Pia in *Nicotiana benthamiana* elicits a weak cell death response

*N. benthamiana* is a well-established model system for assaying the response of rice NLRs to *M. oryzae* effectors (11,12,28). Therefore, we used this system to test whether Pik NLRs would show any response to the effector AVR-Pia. When AVR-Pia was transiently expressed in *N. benthamiana* via agroinfiltration, along with Pikp-1 and Pikp-2, there was a weak cell death response observed, as visualised by a yellowing of the tissue at the infiltration site, and fluorescence under UV light (Fig. 1A). The cell death was weaker compared to AVR-PikD (positive control), but was stronger than for the AVR-PikD point mutant (AVR-PikD^H46E^), a negative control that is not recognised by Pikp (11). To confirm that each protein was expressed, Western blot analysis of extracted leaf tissue was used to assess protein accumulation (Fig. S1). These results show that the Pikp NLRs can respond to AVR-Pia, although the response was limited compared to their ‘matched’ effector AVR-PikD. Interestingly, when the Pikm-1/Pikm-2 pair were tested against the same effectors (AVR-PikD, AVR-PikD^H46E^ and AVR-Pia), there was no response to AVR-Pia in planta (Fig. 1B), despite confirmed expression of all proteins in the leaf tissue (Fig. S1). There was a weak response to the AVR-PikD^H46E^ negative control, as previously observed, due to differences in the AVR-PikD His46 interface with Pikm-HMA compared with Pikp-HMA (12). This suggests that the weak cell death response to AVR-Pia is specific for the Pikp allele.

**Figure 1.**
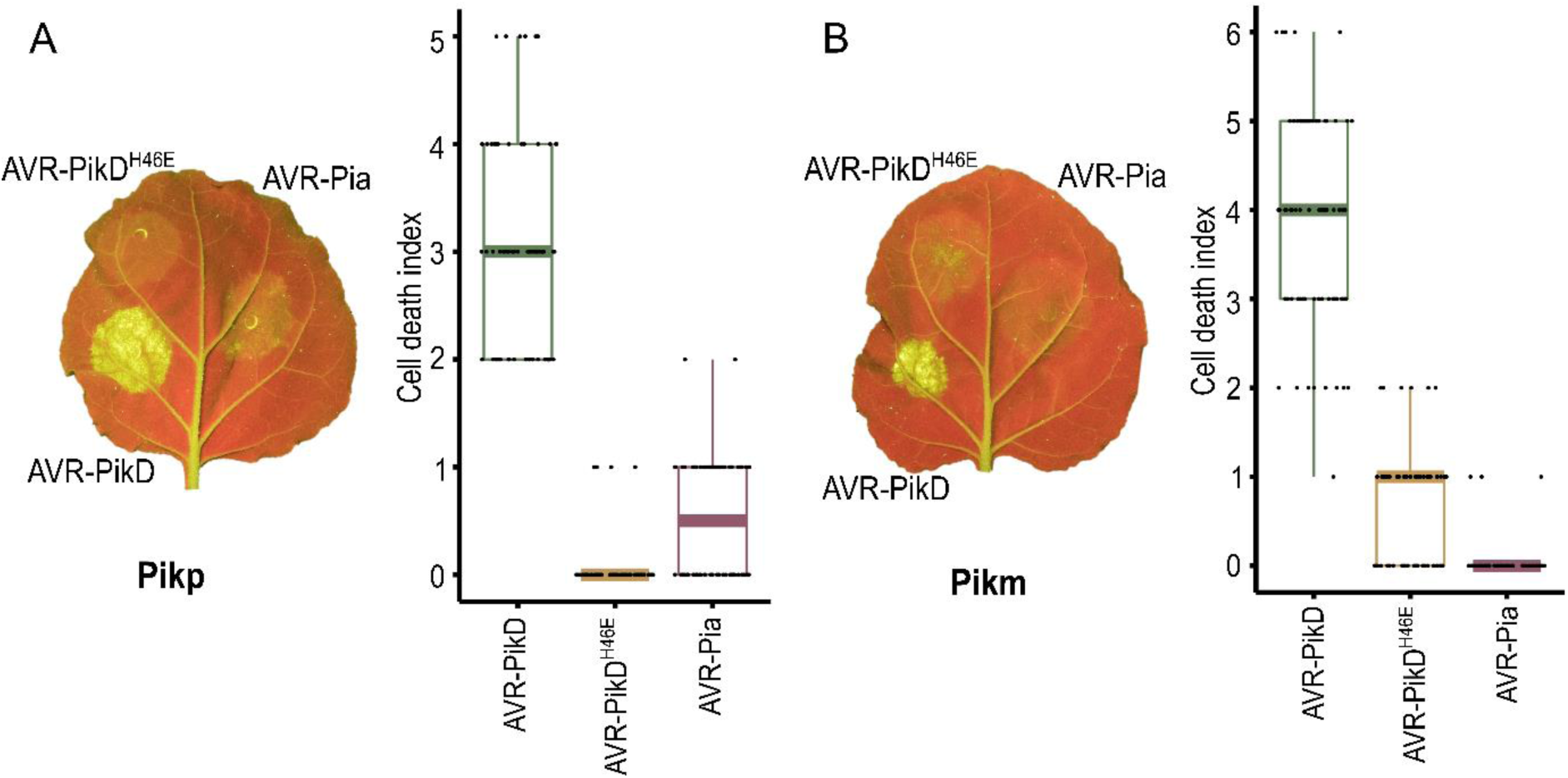
Pikp, but not Pikm, responds weakly to AVR-Pia when transiently expressed in *N. benthamiana.* *N. benthamiana* leaves were visually scored for cell death 5 days post infiltration using the previously published scoring scale (11) from 0 to 6. Representative leaf image shows cell death as autofluorescence under UV light (note: data not used for box plot). Box plots each show 70 repeats of the cell death assay in the example leaf. For each box, the centre line represents the median score, the edges show the upper and lower quartiles and the whiskers show the 1.5x interquartile range. Each data point is represented on the plot as a dot. A) Pikp-1/Pikp-2 transiently expressed with AVR-PikD, AVR-PikD^H46E^ and AVR-Pia. B) Pikm-1/Pikm-2 transiently expressed with AVR-PikD, AVR-PikD^H46E^ and AVR-Pia.

### Rice plants expressing Pikp are partially resistant to *Magnaporthe oryzae* expressing AVR-Pia

Next, we investigated whether the responses of the Pik NLRs to AVR-Pia in *N. benthamiana* could be observed in rice. To test this, we used a spot-inoculation assay to infect rice cultivars with a pathogen strain (Sasa2) transformed to express different effectors. As expected, rice plants that do not express either Pik or RGA NLRs (cv. Nipponbare) are susceptible to infection by all *M. oryzae* Sasa2 lines tested (clear spreading lesions away from the infection site, Fig. 2). Rice plants expressing Pikp (cv. K60) showed resistance to the Sasa2 lines expressing AVR-PikD (positive control) and consistently displayed a reduced susceptibility (partial resistance) phenotype to lines expressing AVR-Pia, developing disease lesions that spread away from the infection site, but are not as developed as the negative controls. This partial resistance phenotype was not observed in rice plants expressing Pikm (cv. Tsuyuake), consistent with results from *N. benthamiana*. Further, rice plants expressing RGA5/RGA4 (cv. Sasanishiki) are susceptible to the Sasa2 line expressing AVR-PikD, showing these NLRs do not partially respond to this effector. All pairwise resistance phenotypes behaved as expected.

**Figure 2.**
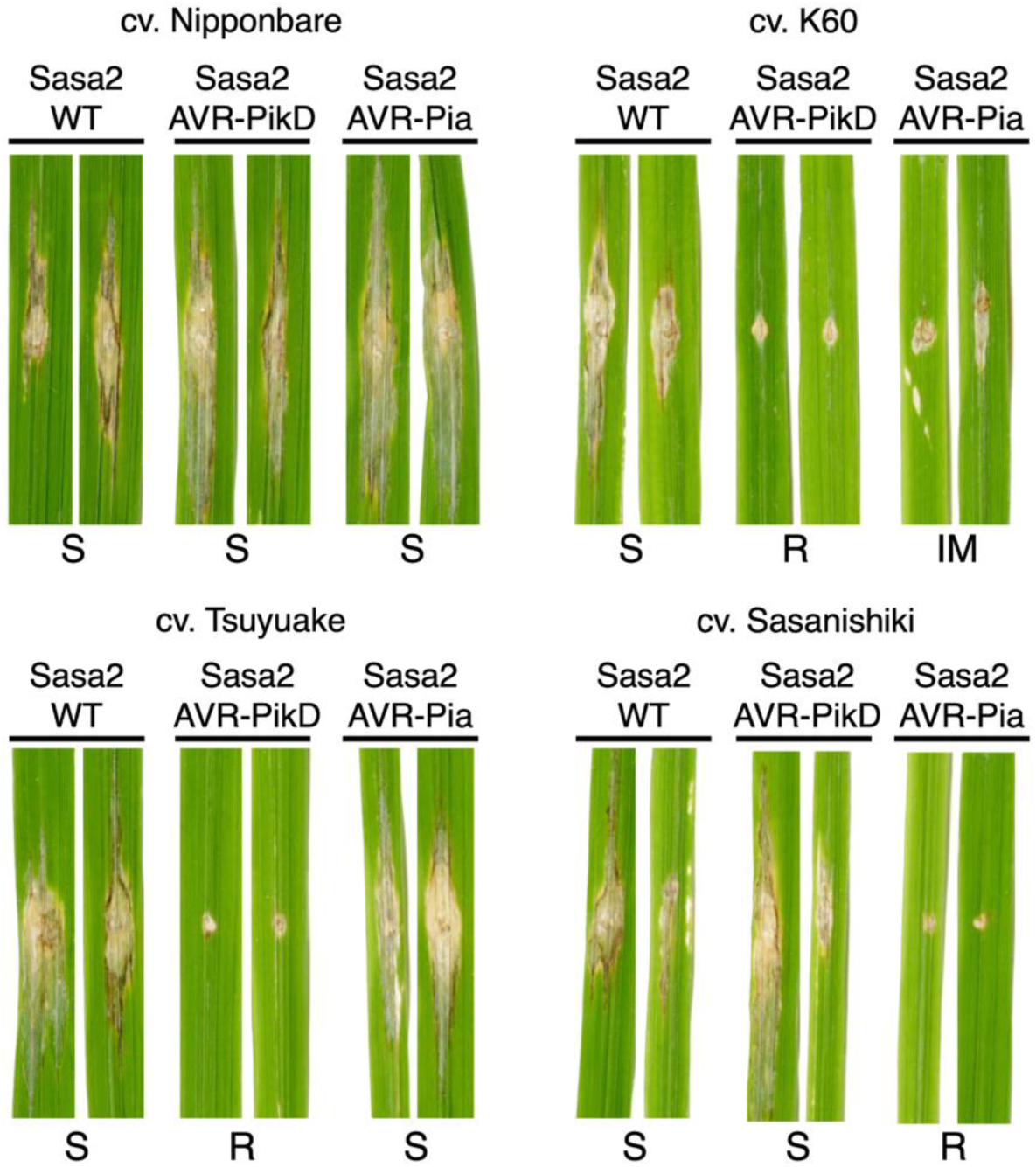
Pikp confers partial resistance to *M. oryzae* expressing AVR-Pia. Images of rice leaves following spot-inoculation assays of Sasa2 *M. oryzae* strain expressing no effectors (WT), AVR-PikD or AVR-Pia,. Strains were inoculated onto rice cultivars containing either Pikp-1/Pikp-2 (cv. K60), Pikm-1/Pikm-2 (cv. Tsuyuake), RGA5/RGA4 (cv. Sasanishiki) or none of the above (cv. Nipponbare). S = Susceptible, R = Resistant, IM = Intermediate. Leaf samples were harvested 10 days post inoculation.

### The HMA domain of Pikp can bind AVR-Pia in vitro

Previously, a tight correlation was observed between in planta response phenotypes in *N. benthamiana* and rice, and in vitro binding between Pik-HMA domains and effectors (12,18). We therefore tested the interaction of Pikp-HMA and Pikm-HMA domains with AVR-Pia following heterologous expression and purification of these proteins.

Firstly, analytical gel filtration was used to qualitatively determine whether Pik-HMA domains and AVR-Pia could form a complex. In isolation, AVR-Pia elutes at a retention volume of 15-15.5 mls (Fig. 3A). When mixed with the Pikm-HMA domain, no change in AVR-Pia retention was observed, consistent with the lack of response in plants. In contrast, when mixed with the Pikp-HMA domain, AVR-Pia elutes earlier at ∼12 mls suggesting a complex is formed, which was confirmed by SDS-PAGE (Fig. S2). Note that Pik-HMA domains do not sufficiently absorb UV light to give a signal in gel filtration under the conditions shown, but can be seen by SDS-PAGE.

**Figure 3.**
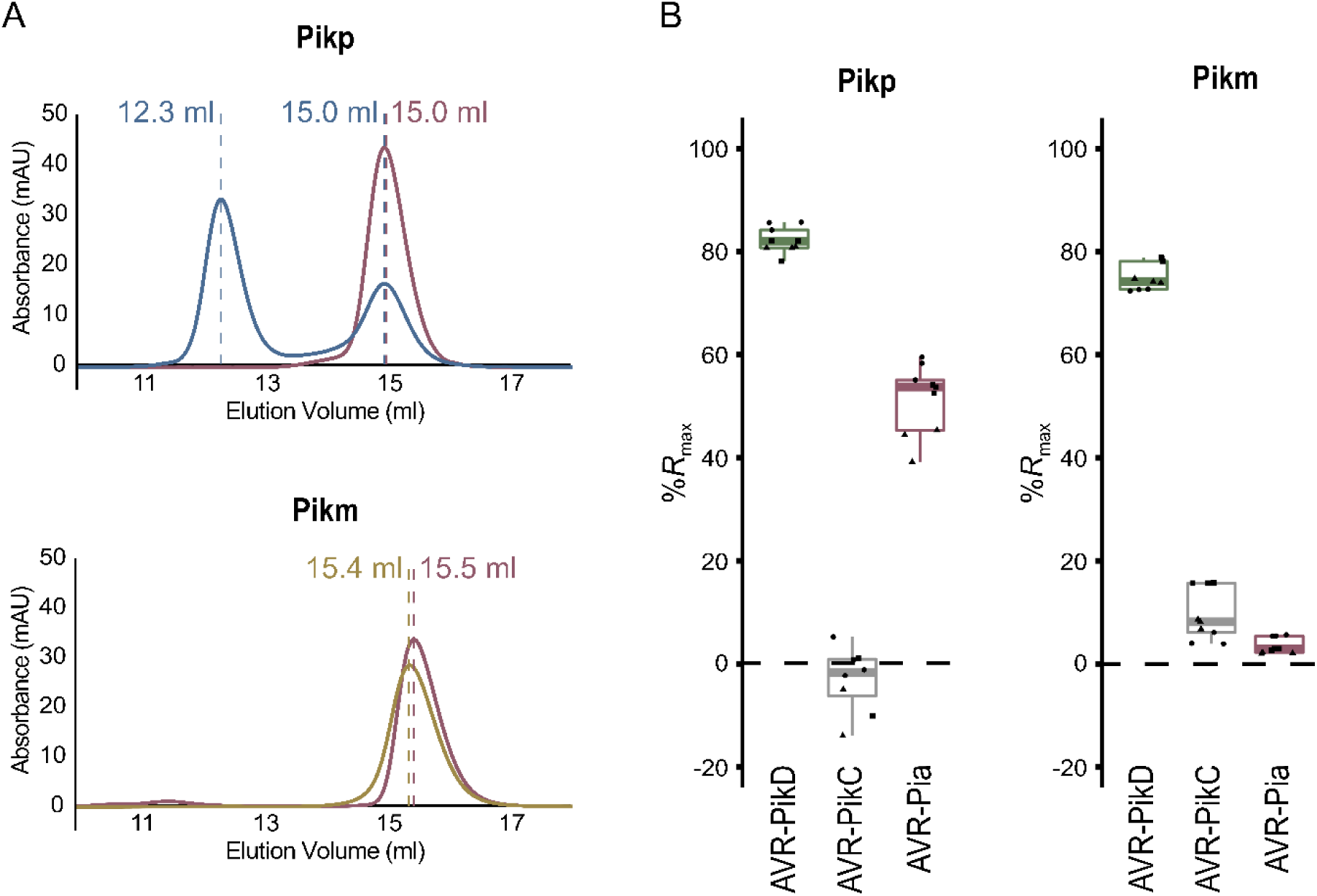
Pikp-HMA, but not Pikm-HMA, binds AVR-Pia in vitro. A) Analytical gel filtration traces assessing complex formation of Pikp-HMA (top panel) and Pikm-HMA (bottom panel) with AVR-Pia. Elution volumes for AVR-Pia alone (pink) and when mixed with Pikp-HMA (blue) and Pikm-HMA (gold) are labelled. Earlier elution indicates a larger molecular mass. SDS-PAGE analysis of eluent at the relevant volumes is shown in Fig. S2. Absorbance observed is only due to the effectors, as Pik-HMA domains essentially do not absorb light at the wavelength measured. B) Surface plasmon resonance data showing *R*_max_ (%) (the percentage of theoretical maximum response for HMA binding to immobilised effector) for Pikp-HMA (left panel) and Pikm-HMA (right panel) at 100 nM concentration binding to AVR-PikD, AVR-PikC or AVR-Pia. Based on previously published data (12), binding was assumed to be 2:1 for Pikp-HMA with AVR-PikD and AVR-PikC, and 1:1 for all other interactions. Box plots show data for three repeats carried out in triplicate, where data points for each repeat are shown as a different shape. Note that only 8 data points are shown for Pikp-HMA with the negative control AVR-PikC, due to poor effector capture in a single run. Equivalent data for 40 nM and 4 nM HMA concentrations are shown in Fig. S2.

We then used surface plasmon resonance (SPR) to measure binding affinities, as described previously (12). These results were expressed as a percentage of the theoretical maximum response (*R*_max_), which gives a relative indication of binding strength. The positive and negative controls for Pikp-HMA and Pikm-HMA binding, the effector variants AVR-PikD and AVR-PikC, show strong and weak/no binding, as expected (Fig. 3B, Fig. S2). Consistent with gel filtration, essentially no binding is observed between Pikm-HMA and AVR-Pia, but Pikp-HMA binds AVR-Pia at ∼50 % *R*_max_ (for the 100 nM Pikp-HMA concentration), independently confirming in vitro interaction and correlating with in planta responses.

### Pikp-HMA binds AVR-Pia at a different interface to AVR-PikD

To visualise the interface formed between Pikp-HMA and AVR-Pia, and compare it to that with AVR-Pik, we purified the complex between these proteins and determined the structure to 1.9 Å resolution using X-ray crystallography. The details of X-ray data collection, structure solution and structure completion are given in Materials and Methods, Table 1 and Fig. S3.

**Table 1:** X-ray data collection and refinement statistics for Pikp-HMA/AVR-Pia.

|  |  |
| --- | --- |
| **Data collection statistics** |  |
| Wavelength (Å) | 0.9763 |
| Space group | *P*22<sub>1</sub>2<sub>1</sub> |
| Cell dimensions |  |
| *a*, *b*, *c* (Å) | 34.84, 53.44, 117.81 |
| $\alpha$, $\beta$, $\gamma$ (°) | 90.00, 90.00, 90.00 |
| Resolution (Å)* | 48.67-1.90 (1.94-1.90) |
| *R*<sub>merge</sub> (%)<sup>#</sup> | 5.7 (122.9) |
| Mean *I*/σ*I*<sup>#</sup> | 19.7 (2.4) |
| Completeness (%)<sup>#</sup> | 100 (100) |
| Unique reflections<sup>#</sup> | 18107 (1151) |
| Redundancy<sup>#</sup> | 12.6 (13.3) |
| CC(1/2) (%)<sup>#</sup> | 99.9 (80.9) |
| **Refinement and model statistics** |  |
| Resolution (Å) | 48.72-1.90 (1.95-1.90) |
| *R*<sub>work</sub>/*R*<sub>free</sub> (%)<sup>^</sup> | 20.3/24.5 (35.8/41.8) |
| No. atoms |  |
| Protein | 2113 |
| Water | 89 |
| Average B-factors (Å<sup>2</sup>) |  |
| Protein | 54.1 |
| Water | 58.1 |
| R.m.s deviations<sup>^</sup> |  |
| Bond lengths (Å) | 0.0117 |
| Bond angles (°) | 1.501 |
| Ramachandran plot (%)** |  |
| Favoured | 98.5 |
| Allowed | 1.5 |
| Outliers | 0 |
| MolProbity Score | 1.52 (95<sup>th</sup> percentile) |

\*The highest resolution shell is shown in parenthesis.
<sup>#</sup>As calculated by Aimless, <sup>^</sup>As calculated by Refmac5, \*\*As calculated by MolProbity

Each partner in the complex adopts a similar overall fold to previously solved structures. Pikp-HMA (11,12) comprises two adjacent α-helices opposite a four-stranded β-sheet (Fig. 4A, B). Previous structures of AVR-Pia were determined by NMR spectroscopy (18,30), and the crystal structure determined here is very similar, comprising the six-stranded β-sandwich characteristic of MAX effectors (18). In the crystal structure, β-5 is not well-defined and appears as a loop joining β-4 and β-6, but overall the configuration of this region is very similar to the NMR ensemble. As previously observed, a disulphide bond is formed between residues Cys25 and Cys66.

**Figure 4.**
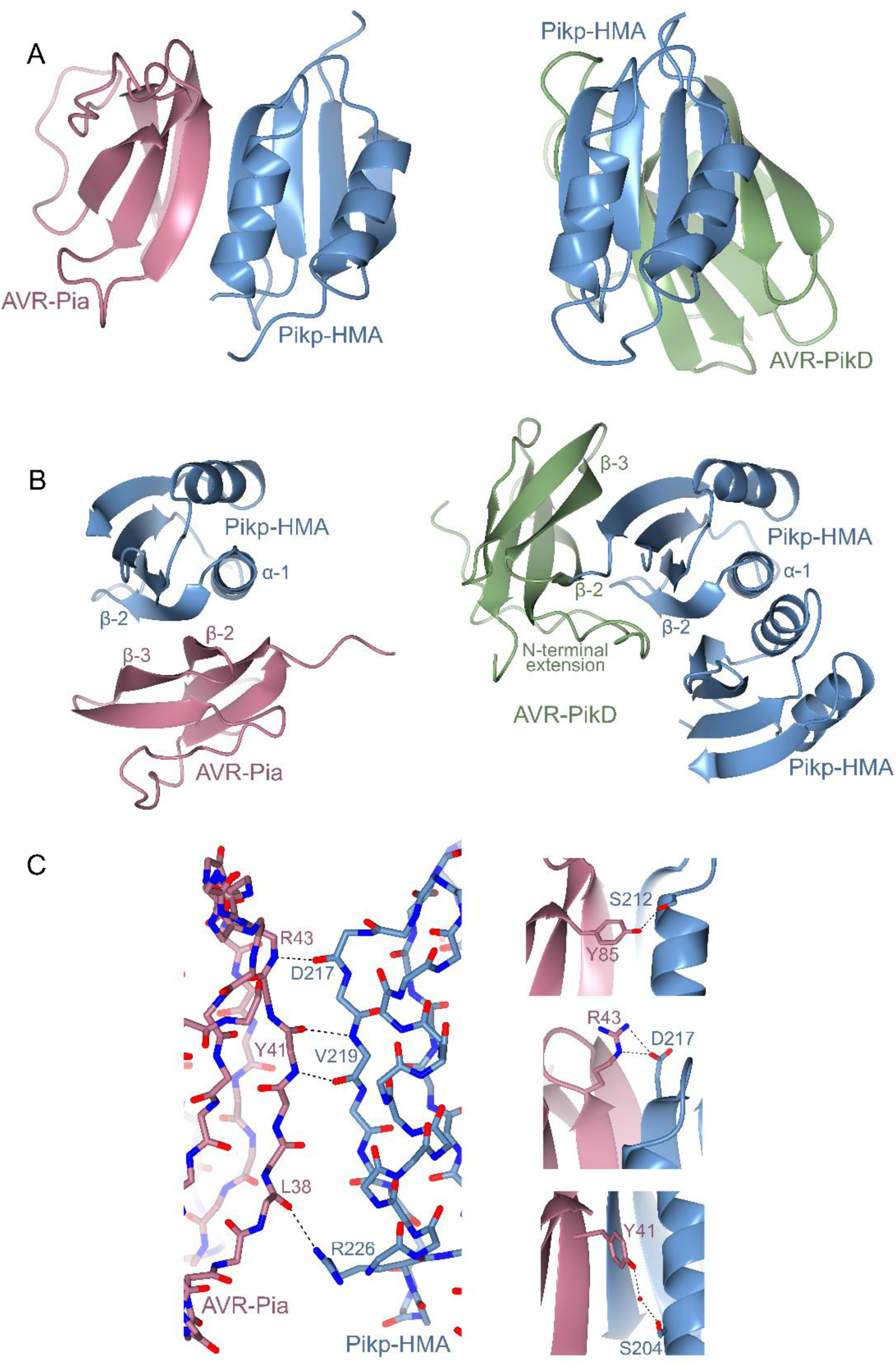
The structural basis of Pikp-HMA interaction with AVR-Pia. A) Schematic diagram of the structure of Pikp-HMA in complex with AVR-Pia refined to 1.9 Å resolution by X-ray crystallography (left), compared to the structure of Pikp-HMA in complex with AVR-PikD (PDB 6G10, right, only a Pikp-HMA monomer displayed here). AVR-Pia is shown in pink, AVR-PikD in green and Pikp-HMA in blue. The Pikp-HMA monomer is shown in the same orientation for both structures. B) An alternative view (rotated ∼90° horizontally and vertically) of the Pikp-HMA/AVR-Pia and Pikp-HMA/AVR-PikD structures shown in A, with secondary structure features labelled (Pikp-HMA dimer structure shown in this view). C) Details of the interface between Pikp-HMA and AVR-Pia, showing interactions at the peptide backbone (left), and selected side-chain interactions (right). Dotted lines show hydrogen bonds, red spheres represent water molecules. Carbons are coloured according to the protein (Pikp-HMA in blue, AVR-Pia in pink) with oxygen atoms shown in red and nitrogen in dark blue. Labels show the single letter amino acid code with position in the peptide chain.

Strikingly, although the two proteins in the complex adopt essentially identical folds to their structures in isolation, Pikp-HMA binds AVR-Pia at a completely different interface to the AVR-Pik effectors (Fig. 4A, B). Whilst Pikp-HMA binds AVR-PikD opposite the face of its β-sheet, it binds AVR-Pia adjacent to α-1 and β-2 (Fig. 4B). In both cases, the positioning of Pikp-HMA relative to the effector allows the formation of a continuous anti-parallel β-sheet between the proteins. In the case of AVR-PikD, the β-strands from Pikp-HMA form a sheet with β-strands 3-5 of AVR-PikD. For AVR-Pia, the β-strands involved are 1, 2 and 6. Another striking feature is that while Pikp-HMA is a dimer in the structure with AVR-PikD (11,12), it is a monomer with AVR-Pia. Indeed, AVR-Pia occupies the same binding surface as the Pikp-HMA dimer in the Pikp-HMA/AVR-PikD structure, which suggests that AVR-Pia binding is competing with Pikp-HMA dimerization in solution.

The interface formed between Pikp-HMA and AVR-Pia covers an area of 460 Å^2^ (as calculated by PISA (31)), approximately half of that seen between Pikp-HMA and AVR-PikD (986 Å^2^ (12)). Further, the interface between Pikp-HMA and AVR-Pia is dominated by hydrogen bonds between the peptide backbone, with the main contributions derived from Pikp-HMA^D217^, Pikp-HMA^V219^, AVR-Pia^Y41^ and AVR-Pia^R43^ (Fig. 4C). The backbone oxygen atom of AVR-Pia^L38^ also forms a hydrogen bond with the side chain of Pikp-HMA^R226^. There are only limited side chain mediated interactions in the Pikp-HMA/AVR-Pia complex, with a hydrogen bond/salt bridge interaction formed between AVR-Pia^R43^ and Pikp^D217^, and the hydroxyl group on the C-terminal residue of AVR-Pia, Tyr85, also forms a hydrogen bond with Pikp^S212^ (Fig. 4C). Finally, an indirect interaction, mediated by a water molecule, is found between the side chains of AVR-Pia^Y41^ and Pikp^S204^ (Fig. 4C). These limited intermolecular interactions and small interface area provide an explanation for the weaker binding affinity seen for Pikp-HMA to AVR-Pia when compared to AVR-PikD in vitro, and reduced responses in planta.

### Pikp recognises AVR-Pia through different molecular features compared to AVR-PikD

Despite only sharing 17 % sequence identity (Fig. S4), AVR-Pia and AVR-PikD both adopt the MAX effector fold. However, AVR-PikD also contains an additional N-terminal extension (comprising residues Arg31 to Pro52) that partially wraps around, and is held in place by, the core structure (see Fig. 4B, Fig. S4). This extension plays a key role in the interaction of AVR-PikD and Pikp-HMA, including a histidine residue (His46), which forms hydrogen bond/salt bridge interactions with Ser218 and Glu230 in Pikp-HMA (11). We considered that modifying the core MAX fold of AVR-Pia to add the AVR-PikD N-terminal extension, might allow Pikp to respond more strongly to the effector by switching the interaction of the chimeric effector (AVR-Pia^NAVR-PikD^) to the ‘AVR-PikD-like’ interface of Pikp-HMA. We also investigated the effect of removing the N-terminal extension from AVR-PikD (AVR-PikD^Δ22-52^).

After generating the appropriate constructs, they were expressed in *N. benthamiana* via agroinfiltration alongside Pikp-1/Pikp-2 or Pikm-1/Pikm-2. In these assays, neither Pikp nor Pikm responded to either AVR-Pia^NAVR-PikD^ or AVR-PikD^Δ22-52^ (Fig. 5). Using western blot analysis, we confirmed the expression of AVR-Pia^NAVR-PikD^ in the infiltrated leaf tissue, suggesting that the lack of cell death is not due to lack of protein accumulation (Fig. S5). However, for AVR-PikD^Δ22-52^, the expression of protein in the leaf tissue is low, suggesting that the N-terminal truncation has destabilised AVR-PikD. These results show that the addition of the N-terminal extension of AVR-PikD to AVR-Pia has not enhanced the response by Pikp or enabled Pikm to respond to the effector. It is possible that the N-terminal extension in AVR-Pia^NAVR-PikD^ remains disordered, and hinders the ability of Pikp to bind to the effector.

**Figure 5.**
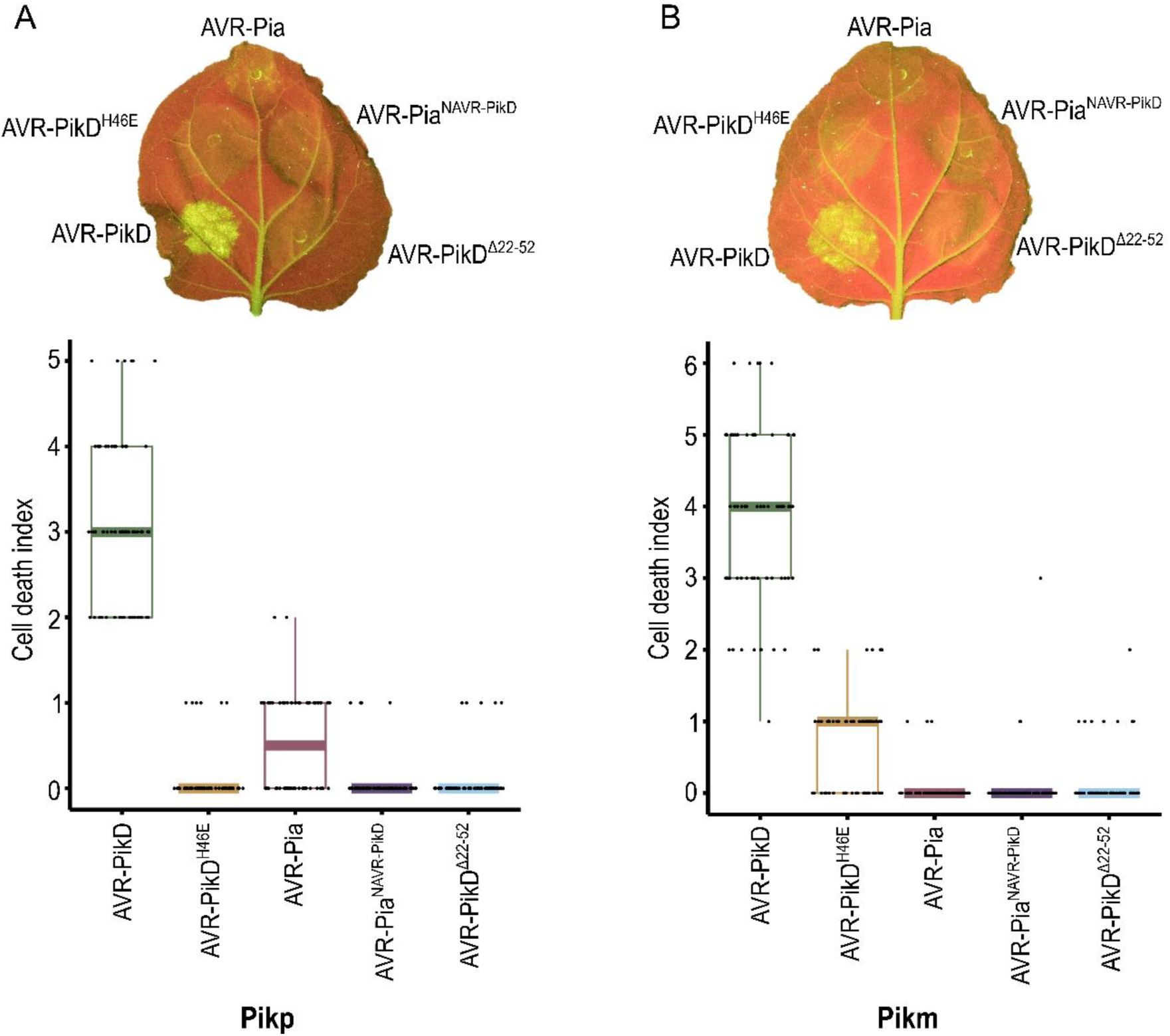
Modifying AVR-Pia with the N-terminal extension of AVR-PikD does not affect the Pik NLR response. *N. benthamiana* leaves were visually scored for cell death 5 days post infiltration using the previously published scoring scale (11) from 0 to 6. Representative leaf image shows cell death as autofluorescence under UV light. Box plots each show 70 repeats of the cell death assay in the example leaf. For each box, the centre line represents the median score, the edges show the upper and lower quartiles and the whiskers show the 1.5x interquartile range. Each data point is represented on the plot as a dot. A) Pikp-1/Pikp-2 transiently expressed with AVR-PikD, AVR-PikD^H46E^, AVR-Pia, AVR-Pia^NAVR-PikD^ and AVR-PikD^Δ22-52^. B) Pikm-1/Pikm-2 transiently expressed with AVR-PikD, AVR-PikD^H46E^, AVR-Pia, AVR-Pia^NAVR-PikD^ and AVR-PikD^Δ22-52^. In A) and B) the data for AVR-PikD, AVR-PikD^H46E^ and AVR-Pia is the same data as shown in Fig. 1, to give direct comparison (all of this data was acquired within the same experimental repeats).

## DISCUSSION

Integrated domains in plant NLR immune receptors bait pathogen effectors to initiate an immune response. Understanding the specificity of effector binding by these integrated domains gives important insights into evolution and function of plant innate immunity. The discovery that rice blast pathogen effectors with a common structural fold can be recognised by the same type of integrated domain in rice NLRs raises questions about specificity, and possible plasticity of recognition. *M. oryzae* MAX effectors AVR-PikD and AVR1-CO39 are bound at different interfaces by their respective NLR-encoded HMA domains (11,12,29). Here, we investigated the interaction of a “mis-matched” NLR integrated domain (Pikp-HMA) and pathogen effector (AVR-Pia), to better understand how protein interfaces contribute to signalling. Ultimately, we hope such studies will lead to improved engineering of NLRs for use in crops.

### A single NLR integrated domain can bait distinct pathogen effectors

Intriguingly, while Pikp-HMA binds AVR-Pia at a different interface to AVR-PikD, it uses the same interface that RGA5-HMA uses to bind AVR1-CO39 (29). Therefore, a single integrated domain in a plant NLR can interact with divergent effectors via different surfaces. Fig. 6 shows a comparison between the Pikp-HMA/AVR-Pia complex and that of the published RGA5-HMA/AVR1-CO39 structure (29) (HMA sequence alignments shown in Fig. S4). Like Pikp-HMA, RGA5-HMA forms a dimer in solution, and binding to the effector competes with this, such that only an HMA monomer is present in each complex (29). Globally, the complexes are very similar, and both rely heavily on peptide backbone interactions for maintaining an interaction between the HMA and effector. One of the most striking differences is the contribution of residues in the N-terminus of AVR1-CO39 (Trp23 and Lys24) to the interaction, which is not seen in the Pikp-HMA/AVR-Pia complex. However, the three important binding regions in the RGA5-HMA/AVR1-CO39 complex noted by Guo et al. are shared by Pikp-HMA/AVR-Pia, although the nature of the residues and interactions involved differ. At the equivalent AVR1-CO39^T41^ and RGA5^D1026^ binding area, there is a side chain interaction between AVR-Pia^R43^ and Pikp^D217^. At an equivalent location to the second binding area (AVR1-CO39^I39^ and RGA5^V1028^), there are AVR-Pia^Y41^ and Pikp^V219^ backbone interactions and a water-mediated hydrogen bond between the side chain of AVR-Pia^Y41^ and Pikp^S204^. Finally, the third binding area involves a backbone interaction between Ile1030 of RGA5-HMA and Asn37 of AVR1-CO39. At a similar area in the Pikp-HMA/AVR-Pia interface, there is a hydrogen bond between the backbone of AVR-Pia^L38^ and the side chain of Pikp^R226^. The overall close similarities between these complexes imply that this is a biologically relevant interface and suggest that AVR-Pia may interact with RGA5-HMA at the same interface, although there is no structural confirmation of this to date.

**Figure 6.**
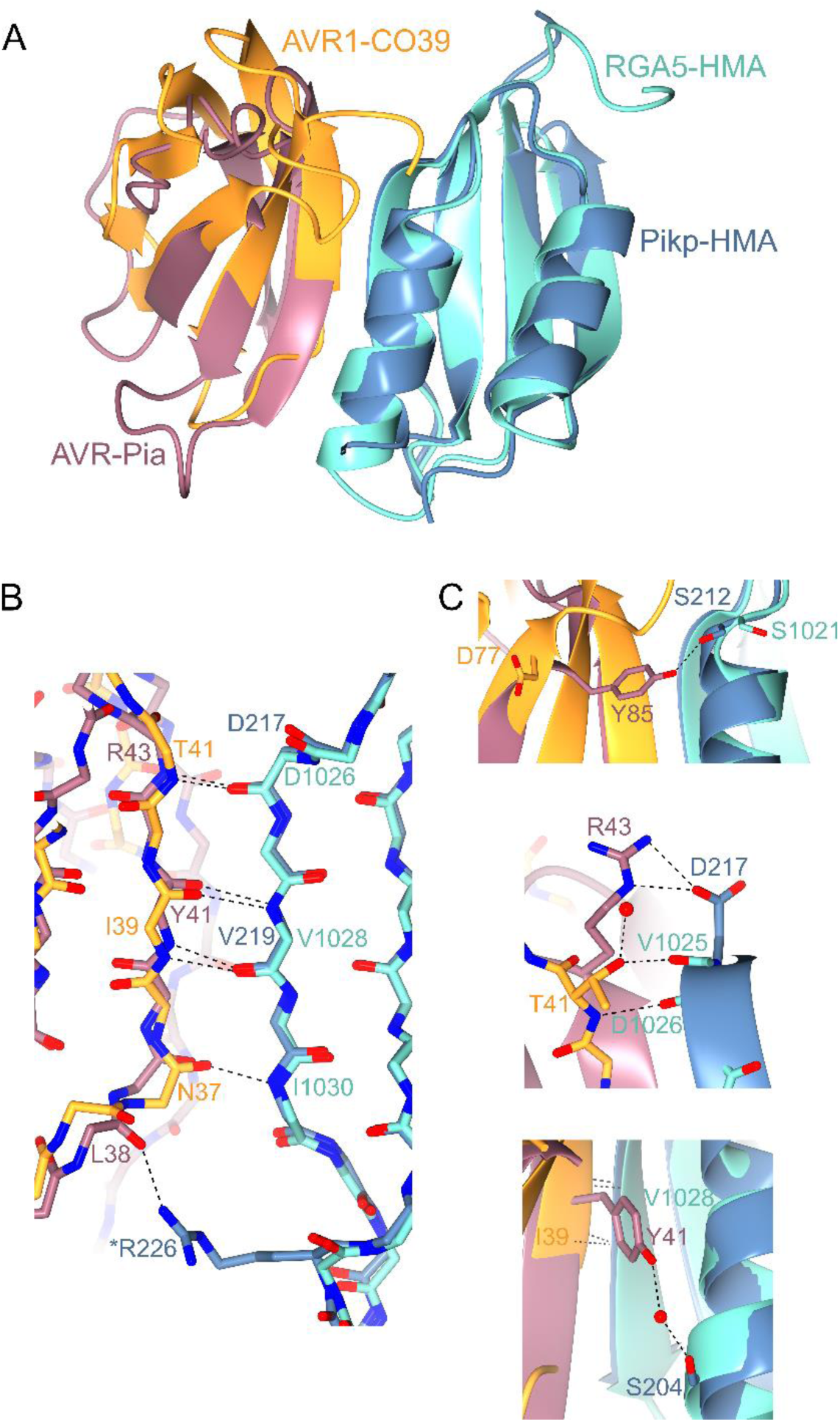
Structural comparison of Pikp-HMA/AVR-Pia and RGA5-HMA/AVR1-CO39 complexes. Overlays of Pikp-HMA/AVR-Pia with RGA5-HMA/AVR1-CO39 (PDB 5ZNG), superposed on the HMA domain (RMSD 0.81 Å over 73 residues). AVR-Pia is shown in pink, Pikp-HMA in blue, AVR1-CO39 in orange and RGA5-HMA in turquoise. A) Cartoon ribbon structure to represent overall structures. B) Details of interactions between the peptide backbones at the interface. Dotted lines show hydrogen bonds, carbons are coloured according to the chain with oxygen atoms shown in red and nitrogen in dark blue. Labels show the single letter amino acid code (coloured according to protein) with position in the peptide chain. ‘*’ Indicates a side chain, rather than backbone interaction. C) Further details of important interactions at the interfaces. Red spheres represent water molecules.

While the different HMA domains of RGA5 and Pik use different interfaces to interact with their cognate effectors, Pikp has the capacity to use both of these for binding different effectors. Our initial observations in rice suggest that RGA5/RGA4 cannot respond to AVR-PikD, indicating that RGA5 might not be able to use the alternative “AVR-PikD-like” binding interface. We hypothesise that following HMA domain integration into their respective ancestor proteins, Pik-1 and RGA5 have evolved to respond to their cognate effectors through variation both within the HMA domains, but also within the rest of the NLR architecture. The position of the HMA domain integration is likely critical, and may affect available HMA-binding interfaces for both the effectors and intra-/inter-molecular interactions within the NLRs that support downstream signalling.

### The Pik NLR response to and interaction with AVR-Pia is allele-specific

Pikm is not able to respond to AVR-Pia, despite both Pikp and Pikm recognising the same MAX effector AVR-PikD. When the structure of the Pikp-HMA/AVR-Pia complex is overlaid with Pikm-HMA (12), the overall HMA conformation is nearly identical, but sequence diversity results in different side chains being presented at the predicted interaction surface. Most apparent is that Pikp-HMA^D217^, which forms a hydrogen bond/salt bridge interaction with AVR-Pia^R43^, is replaced by a histidine residue at the equivalent position in Pikm-HMA. This change may, in part, account for a reduced affinity for AVR-Pia, although it seems unlikely to fully account for a lack of interaction. Further experiments are required to investigate why Pikm-HMA does not bind AVR-Pia in vitro or Pikm respond to AVR-Pia in planta.

### Using integrated domain cross-reactivity for NLR engineering

The cross-reactivity of Pikp for the “mis-matched” AVR-Pia effector raises exciting possibilities around engineering Pikp to respond more robustly to this effector, whilst maintaining AVR-PikD interactions. As noted by Guo et al., the use of different interfaces for the effectors may allow engineering of one surface without significantly disrupting the binding at the other (29). Such detailed structural knowledge paves the way towards future NLR engineering for improved disease resistance that may be applicable to other NLR/effector pairs.

## EXPERIMENTAL PROCEDURES

### Cloning and construct generation

Constructs for *N. benthamiana* cell death assays were generated by Golden Gate cloning methods (32). Domesticated Pik-1 and Pik-2 NLRs were used as described in de la Concepcion (12) and each effector construct was generated with an N-terminal 4xMyc tag, a Ubi10 promoter (from *A. thaliana*) and 35S terminator.

For in vitro studies, isolated Pikp-HMA (residues 186-263) and Pikm-HMA (residues 186-264) domain constructs were used as described in de la Concepcion (12). For analytical gel filtration and crystallography studies, AVR-Pia (residues 20-85) was cloned into the pOPINS3C vector by In-Fusion cloning (33) to yield a cleavable N-terminal 6xHis-SUMO tagged construct. For surface plasmon resonance, effectors were amplified from pOPINS3C and cloned into pOPINE to yield a non-cleavable C-terminal 6xHis tag in addition to the SUMO tag, following the strategy used in (11).

### *N. benthamiana* cell death assays

Transient in planta expression, cell death assays and confirmation of protein expression was carried out as described by de la Concepcion et al. (12). Briefly, *Agrobacterium tumefaciens* GV3101 was used to deliver T-DNA constructs into 4-week-old *N. benthamiana* plants (grown at high light intensity, 22-25 °C). Pik-1, Pik-2, AVR-Pik and the P19 suppressor of silencing were mixed prior to infiltration and delivered at OD_600_ 0.4, 0.4, 0.6 and 0.1 respectively. At 5 dpi, detached leaves were imaged under UV light on the abaxial side, and visually scored against a cell death index described previously (11). Scores for three independent repeats are shown as box and whisker plots, generated using R (34) and graphics package ggplot2 (35). Leaf disks taken from representative infiltration spots were frozen, ground and mixed with 2x w/v extraction buffer (25 mM Tris, pH 7.5, 150 mM NaCl, 1 mM EDTA, 10 % v/v glycerol, 10 mM DTT, 2 % w/v PVPP, 0.1 % Tween®-20, 1x plant protease inhibitor cocktail (Sigma)), before centrifugation, filtering and SDS-PAGE/Western blot analysis.

### Rice pathogenicity assays

*M. oryzae* strains Sasa2 and Sasa2 expressing *AVR-PikD* (the transformant harboring 22p:pex31-D (*AVR-PikD* allele fused with the promoter region of *AVR-Pia*)) used in this study are stored at the Iwate Biotechnology Research Center (21). To obtain protoplasts, hyphae of Sasa2 strain were incubated for 3 days in 200 ml of YG medium (0.5% yeast extract and 2% glucose, wt/vol). Protoplast preparation and transformation with pex22p:pex22 (*AVR-Pia* fused with the promoter region of *AVR-Pia*) were performed as previously described (36) to generate Sasa2 strain expressing *AVR-Pia*. Bialaphos-resistant transformants were selected on plates with 250 μg/ml of Bialaphos (Wako Pure Chemicals).

Rice leaf blade spot inoculations were performed with *M. oryzae* strains as previously described (37). Disease lesions were scanned 14 days post-inoculation (dpi). The assays were repeated at least 3 times with similar results.

### Expression and purification of proteins for in vitro studies

All proteins for in vitro studies were expressed from *E. coli* SHuffle cells (38) in auto-induction media (39). Cell cultures were grown at 30 °C for 5 hours, followed by 16 °C overnight. Proteins were purified as described in Maqbool et al. (11). In outline, pelleted cells were lysed by sonication and clarified by centrifugation, before tandem Ni-IMAC and gel filtration protein purification steps using an ÄKTAxpress (GE Healthcare) system. Overnight cleavage of solubility tags with 3C protease at 4 oC, followed by solubility tag removal and a further cycle of gel filtration, yielded purified protein that was concentrated by ultrafiltration and stored at −80° C.

### Expression and purification of proteins for crystallisation

To prepare the Pikp-HMA/AVR-Pia complex for crystallisation studies, separate cell cultures of SUMO-tagged AVR-Pia and 6xHis-MBP-tagged Pikp-HMA were grown and harvested as described previously. After initial protein purification and immediately following removal of the solubility tags, both proteins were combined and subsequently treated as a single sample for the final gel filtration purification stage.

### Protein-protein interaction studies in vitro

Analytical gel filtration and surface plasmon resonance experiments were carried out as described in de la Concepcion et al. (12). For analytical gel filtration, purified proteins were run down a Superdex™ 75 10/300 column (GE Healthcare) at 0.5 ml/min either alone or mixed to assess complex formation (mixtures were incubated on ice for 2 hours prior to experiment). Effectors were used at 50 µM final concentration, and Pikp-HMA and Pikm-HMA were used at 100 µM and 50 µM respectively, to account for dimer formation in solution. For surface plasmon resonance experiments, the Biacore T200 system (GE Healthcare) was used, and C-terminal 6xHis-tagged effector proteins were immobilised onto a NTA sensor chip (GE Healthcare). HMA protein was flowed over the immobilised effector at 30 µl/min (360 sec contact time and 180 sec dissociation time) at 4, 40 and 100 nM concentrations, considering HMA dimer formation where appropriate. The response of a reference cell was subtracted for each measurement. Raw data was exported, % *R*_max_ values were calculated in Microsoft Excel, and then individual % *R*_max_ data from three separate experiments were displayed as box plots in R.

### Crystallisation, data collection and structure determination

For crystallisation, Pikp-HMA/AVR-Pia complex (in a buffer of 20 mM HEPES, 150 mM NaCl, pH 7.5) was used in sitting drop vapour diffusion experiments. Drops were set up in 96-well plates, composed of 0.3 µl purified protein with 0.3 µl reservoir solution, dispensed using the Oryx Nano crystallisation robot (Douglas Instruments). Crystals for data collection were obtained in the Morpheus® screen (Molecular Dimensions), using protein at 18 mg/ml (measured by Direct Detect® spectrometer (Merck)). The crystals were found in well D2 of the screen, and the conditions in this well were: 0.12 M Alcohols (0.2 M 1,6-Hexanediol; 0.2 M 1-Butanol; 0.2 M 1,2-Propanediol; 0.2 M 2-Propanol; 0.2 M 1,4-Butanediol; 0.2 M 1,3-Propanediol), 0.1 M Buffer System 1 (1.0 M imidazole; MES monohydrate (acid), pH 6.5) and 50 % v/v Precipitant Mix 2 (40 % v/v Ethylene glycol; 20 % w/v PEG 8000). Crystals were frozen in liquid nitrogen and X-ray data were collected at the Diamond Light Source (Oxfordshire) on beamline DLS-i03. Crystallographic data was processed using the Xia2 pipeline (40) and AIMLESS (41), as implemented in the CCP4 software suite (42). To solve the structure, a single model from the ensemble of AVR-Pia (PDB file 2MYW) and a monomer structure of Pikp-HMA (PDB accession 5A6P) were used for molecular replacement in PHASER (43). COOT (44) was used for manual rebuilding, and successive rounds of manual rebuilding were followed by rounds of refinement using REFMAC5 (45). The structure was validated using tools provided in COOT, and finally assessed by MolProbity (46). All structure figures were prepared using the CCP4 molecular graphics program (CCP4MG) (42).

### Supporting Information

Supplementary Figures 1 – 5 are shown in the Supporting Information.

### Accession codes

The protein structure of the complex between Pikp-HMA and AVR-Pia, and the data used to derive this, have been deposited at the PDB with accession number 6Q76.

## Acknowledgements

We thank the Diamond Light Source (beamline i03, under proposal MX13467) for access to X-ray data collection facilities. We also thank D. Lawson and C. Stevenson (JIC X-ray Crystallography/Biophysical Analysis Platform) for help with X-ray data collection and SPR. This work was supported by: Biotechnology and Biological Sciences Research Council (BBSRC, UK), grant nos. BB/P012574, BB/M02198X; the ERC (proposal 743165), the John Innes Foundation, the Gatsby Charitable Foundation and JSPS KAKENHI 15H05779.

