## Supplemental Information for "Cross-reactivity of a rice NLR immune receptor to distinct effectors from the blast pathogen leads to partial disease resistance"

### Supporting Information

### Figures S1 - S5

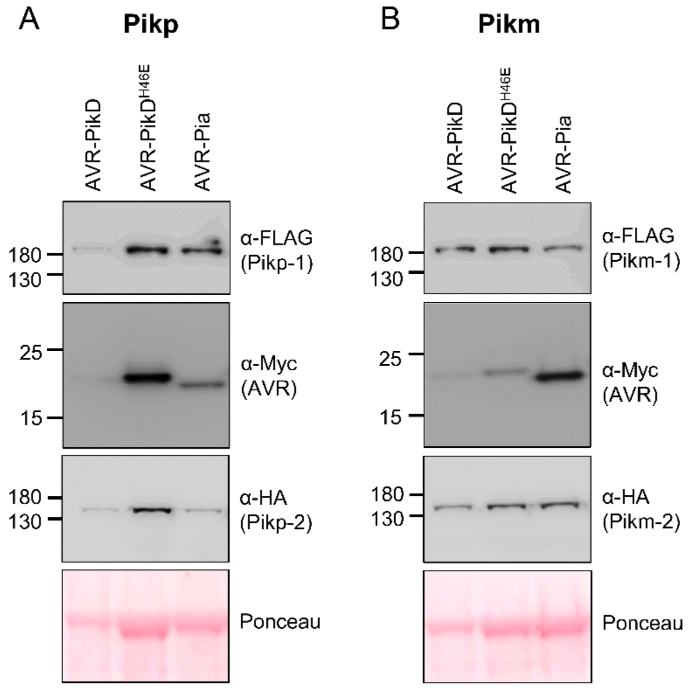


**Figure S1.** Western blots showing protein accumulation following transient expression in *N. benthamiana*, 5 days post agroinfiltration. Blot is representative of 3 biological repeats giving similar results. A) Pikp-1/Pikp-2 with AVR-PikD, AVR-PikD$^{H46E}$ and AVR-Pia. B) Pikm-1/Pikm-2 with AVR-PikD, AVR-PikD$^{H46E}$ and AVR-Pia. Note: the amount of each protein in the Pik-1/Pik-2/AVR-PikD sample appears lower than the others (as indicated in the Ponceau image for total loading) due to greater cell death in this sample, limiting accumulation.

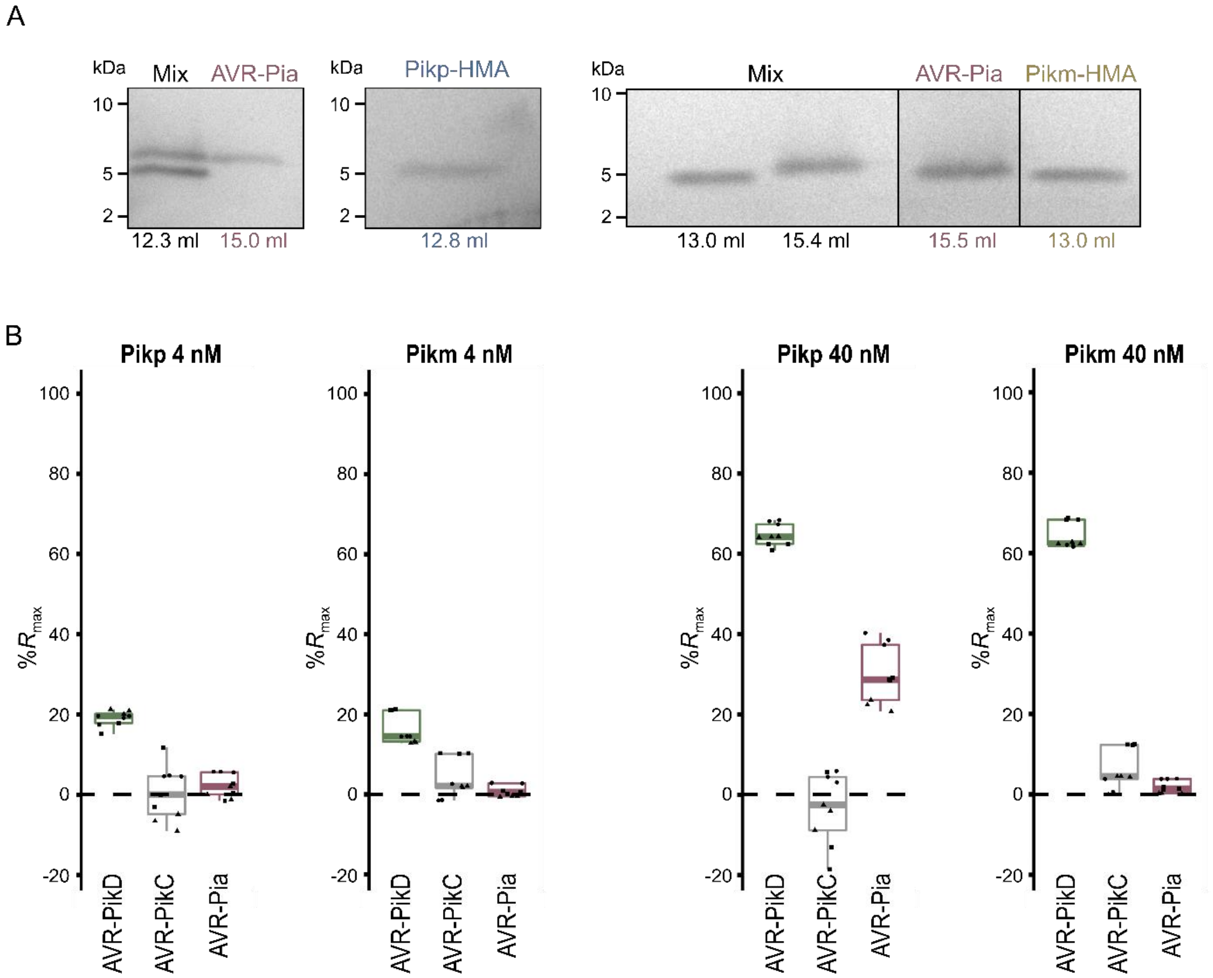


**Figure S2.** A) SDS-PAGE analysis showing the proteins eluted at the volumes indicated in the analytical gel filtration traces shown in Fig. 3A. The isolated HMA domains are not shown in Fig. 3A because they do not absorb sufficient UV light to be detected. B) Surface plasmon resonance $R_{max}$ (%) data for Pikp-HMA and Pikm-HMA at 4 nM and 40 nM concentrations binding to AVR-PikD, AVR-PikC and AVR-Pia. Data is displayed in the same manner as shown in Fig. 3B.

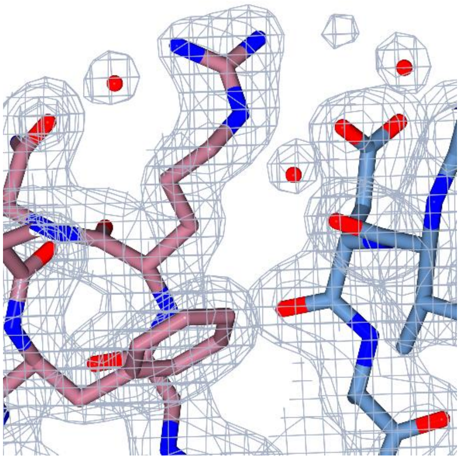

**Figure S3.** Image showing the Pikp-HMA/AVR-Pia structure modelled into electron density (shown in grey mesh), at an interface region between the two proteins (centred around AVR-Pia$^{R43}$ and Pikp-HMA$^{D217}$). Atoms are coloured as described in Fig. 4.

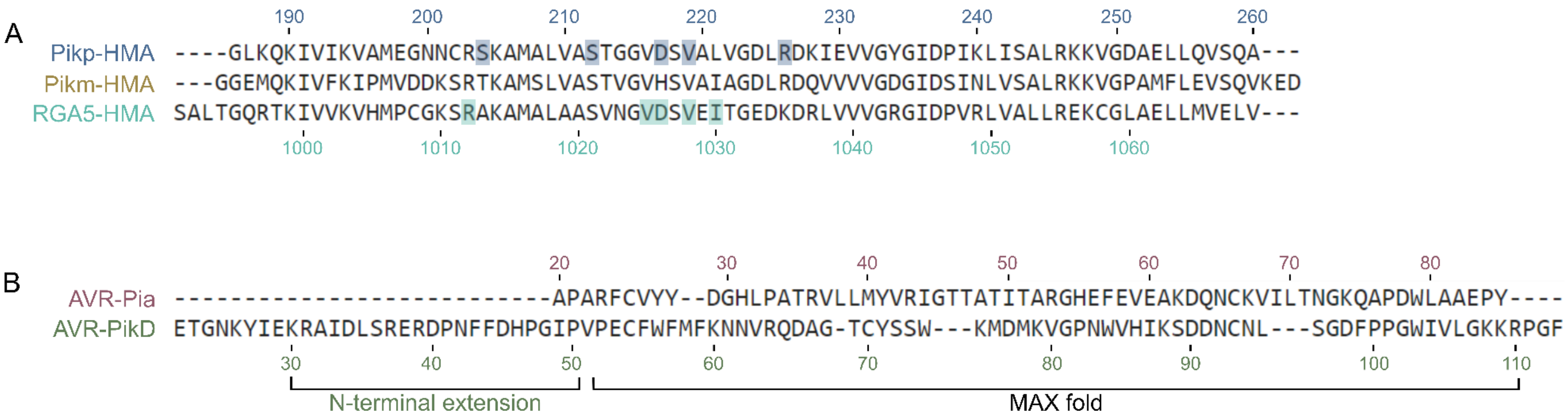


**Figure S4.** A) Sequence alignment of Pikp-HMA, Pikm-HMA and RGA5-HMA, showing only residues present in the PDB accessions 6Q76, 6FU9 (1) and 5ZNG (2) respectively. Residues are numbered for Pikp-HMA and RGA5-HMA. Some key residues for the interaction with AVR-Pia or AVR1-CO39 (as described in the text and in (2)) are highlighted. B) Sequence alignment of AVR-Pia and AVR-PikD, with signal peptides removed. Areas comprising MAX fold (for both effectors) and N-terminal extension (for AVR-PikD only) are indicated.

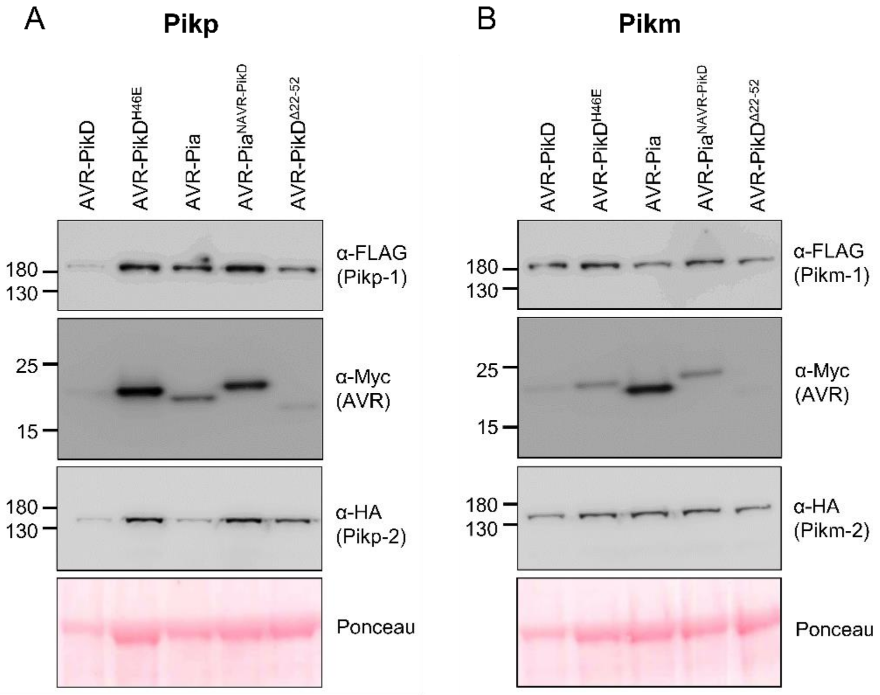


**Figure S5.** Western blots showing protein accumulation following transient expression in *N. benthamiana*, 5 days post agroinfiltration. Blot is representative of 3 biological repeats giving similar results. A) Pikp-1/Pikp-2 with AVR-PikD, AVR-PikD$^{H46E}$, AVR-Pia, AVR-Pia$^{NAVR\text{-}PikD}$ and AVR-PikD$^{\Delta 22\text{-}52}$. B) Pikm-1/Pikm-2 with AVR-PikD, AVR-PikD$^{H46E}$, AVR-Pia, AVR-Pia$^{NAVR\text{-}PikD}$ and AVR-PikD$^{\Delta 22\text{-}52}$. Note: A cropped image of this blot is shown in Fig. S1. All samples were analysed together for direct comparison.
